# The distribution of large floating seagrass (*Zostera marina*) clumps in northern temperate zones of Bohai Bay in the Bohai Sea, China

**DOI:** 10.1101/372565

**Authors:** Xu Min, Zhou Yi, Zhang Tao, Zhang Yun-Ling

## Abstract

Seagrass meadows (*Zostera marina*) are important coastal ecosystems with high levels of productivity and biodiversity. They are subject to considerable natural and anthropogenic threats in China, such as oyster and snail aquaculture, wastewater discharge, electro-fishing, shellfish collection, typhoons and floods. When seagrass communities are disturbed, they can become removed from the sediment and converted into floating clumps, which then serve as marine hot spots attracting a variety of marine organisms that then inhabit them. They are important nursery habitats for many economic fish such as red drum (*Sciaenops ocellatus*), Atlantic cod (*Gadus morhua*), queen conch (*Strombus gigas*), and blue crab (*Callinectes sapidus*). Thus, it is necessary to study the distribution and biological characteristics of these floating seagrass clumps. In September 2016 we observed large scale floating *Z. marina clumps* in the northernmost area of Bohai Bay (38°57’1.14”−39° 0’41.28” N, 118°45’23.22”−118°47’6.96” E), in the Bohai Sea, China. We observed characteristics that precluded their origination from the nearby Caofeidian seagrass meadows. Two research cruises were undertaken, during which we did not observe other marine organisms accompanying these floating *Z. marina* clumps. The dominant frond lengths were 40–50 cm, with less than 5% of the total number of fronds found in larger size categories (80–90 and 90–100 cm). We aim to pursue future research into the breakdown and dislodgement characteristics of *Z. marina* clumps and the processes whereby they sink and integrate with the sediment.

## Introduction

Seagrass meadows are highly biodiverse and productive coastal ecosystems, covering approximately 0.15% of the global ocean area and contributing 1% of the marine net primary production [1]. Seagrass meadows are important nursery habitats for many economically valuable fish and invertebrate species such as red drum (*Sciaenops ocellatus*), Atlantic cod (*Gadus morhua*), queen conch (*Strombus gigas*), and blue crab (*Callinectes sapidus*) [1-2]. They also provide suitable substrate for a variety of marine epiphytes and facilitate the sedimentation of suspended particulates [3-4]. In the past half century, seagrass meadows have declined rapidly worldwide. For example, approximately 2.6 × 10^4^ km^2^ of seagrass meadows, accounting for 15% of the estimated global total, disappeared between 1993 and 2003 [5].

Floating seagrass clumps are an important method of transport for seagrass detritus from one habitat to another [6]. As seagrass meadows age, senesce and interact physically with the environment around them, leaves may break off from the beds and either sink to the surrounding seafloor or float to the sea surface [6]. Due to the high primary productivity of seagrass meadows [7], considerable volumes of leaves are continually shed and transported away from the beds by currents and wave action [8-10]. Large aggregations of this floating vegetation called “seagrass wrack” can form [11] and the biomass is ultimately exported to the seafloor or washed ashore on beaches.

Floating at the sea surface, seagrass wracks can serve as habitat “hot spots” similar to other floating vegetation such as floating macroalgae (e.g. *Sargassum* spp.). Thresher et al.[12] is one of the only studies to report on the trophic impacts of floating seagrass wracks. Through indirect lines of evidence, they reported that microbial decomposition of floating seagrass played a pivotal role in the coastal planktonic food chain [12].

A broad seagrass meadow (39°00’N−39°05’N, 118°41’E−118°44’E), covering approximately 10 km^2^ was discovered in the northwestern area of Longdao Island, Caofeidian, Bohai in October 2015 [13]. To date, it is the largest seagrass meadow discovered in the Bohai Sea and Yellow Sea areas in China, and its composition is dominated by *Zostera marina* [13]. *Z. marina* is a species of seagrass known by the common names ‘common eelgrass’ and ‘seawrack’, and is widely distributed in the Northern Hemisphere. The plant is a rhizomatous herb which produces a long stem with hair-like green leaves. The rhizome grows horizontally through the substrate, anchoring via clusters of roots at nodes. In September 2016, a large area of floating *Z. marina* clumps was observed in the area of Xiang-yun Island, the northernmost area of Bohai Bay in the Bohai Sea, China, nearby to the Caofeidian area. We conducted two research cruises, a week apart, to survey the distribution of these floating *Z. marina* clumps in September 2016. The present study is the first to report the distribution of floating *Z. marina* clumps in temperate zones in the Bohai Sea, China.

## Materials and methods

The field survey was approved by Institute of Oceanology, Chinese Academy of Sciences.

A survey of the distribution of floating *Z. marina* around Xiang-yun island, the northernmost part of Bohai Bay, Bohai Sea was carried out using transect lines at a distance of approximately 0.5–1.0 km (Fig. 1). Sightings of drifting clumps are difficult unless the sea is calm, and only clumps within 50 to 100 m of the vessel were observed. The first cruise was made on 6 September 2016 from ‘Point 1’ to ‘Point 8’, while the second cruise was made on 13 September 2016 from ‘Point 8’ to ‘Point 15’ (Fig. 1). The research vessel kept a stable speed of six knots, and seagrass clumps seen within each transect line were quantified as follows: none, frequent clumps and large amounts (Fig. 2).

**Fig. 1.**
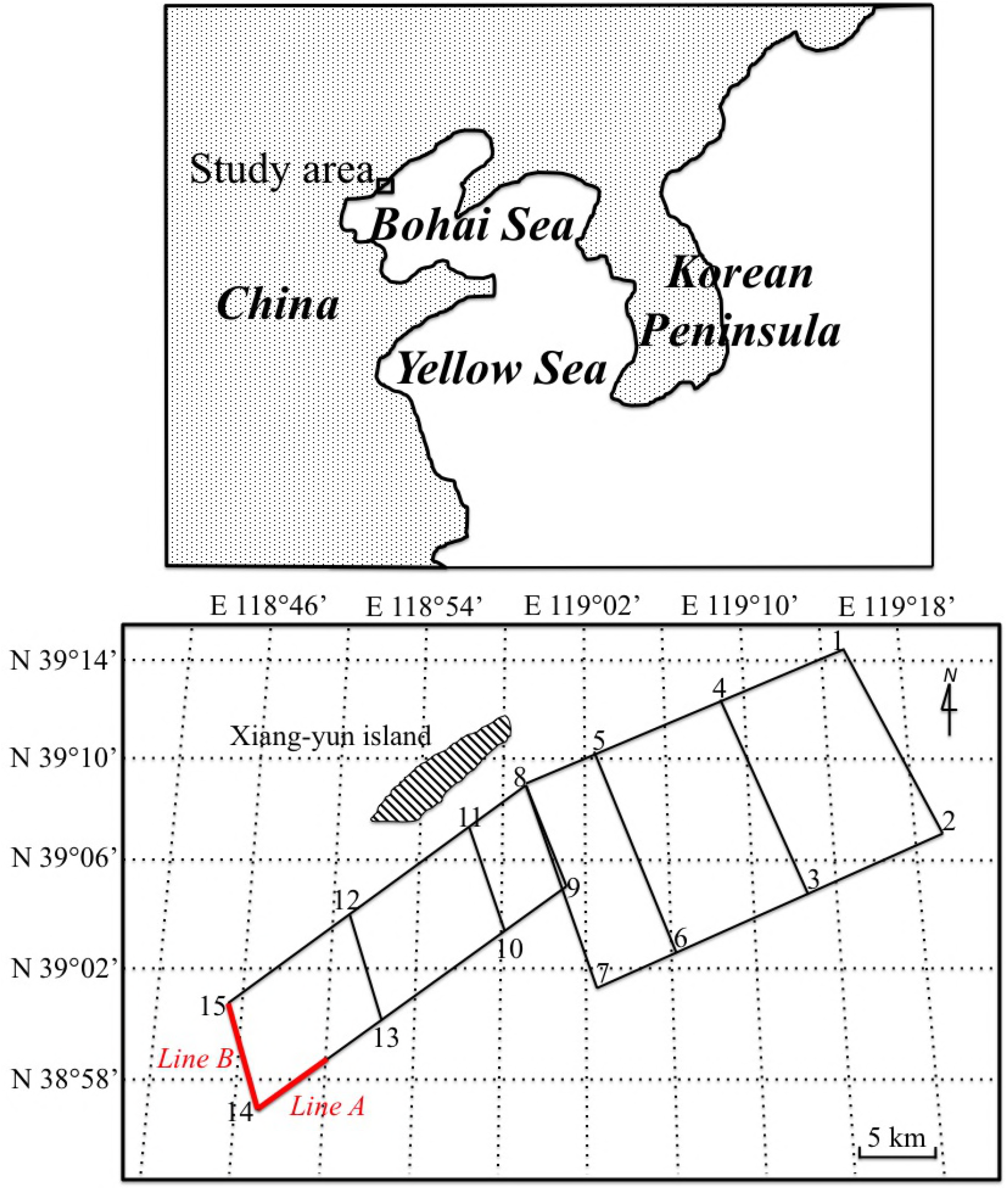
Schematic map showing the route of two research cruises to survey the distribution of floating *Zostera marina* clumps in the waters off Xiang-yun Island, Tangshan in the northernmost area of Baihai Bay in the Bohai Sea, China. On 6 and 13 September 2016 the vessel travelled from ‘Point 1’ to ‘Point 8’ and from ‘Point 8’ to ‘Point 15’, respectively. The red lines (‘Line A’ and ‘Line B’) indicate the locations of floating *Z. marina* clumps. Line A was from N 38°58.9358’ E 118°50.1150’ to ‘Point 14’, and Line B was from ‘Point 14’ to ‘Point 15’. The Caofeidian seagrass meadows are located approximately 6,400 m away from Point 14, 10,000 m away from Point 15 and 13,500m away from N 38°58.9358’ E 118°50.1150’.

**Fig. 2.**
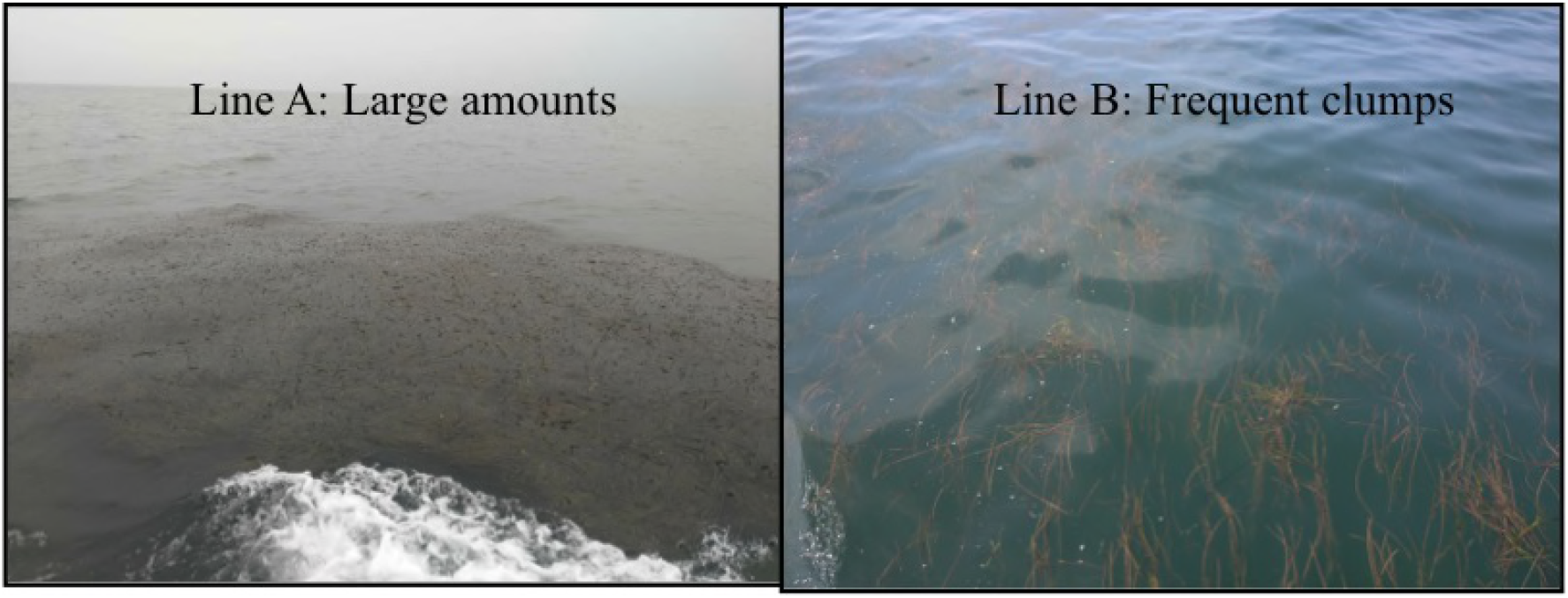
Left: An example of ‘large amounts’ of *Zostera marina* observed in ‘Line A’. Right: An example of floating *Z. marina* leaves observed in ‘Line B’.

Based on previous literature [14-15], a dip net with a ring diameter of 50 cm and mesh size of 5.0 cm was selected for optimal sampling. The net was dipped under each clump, hauled up in one quick sweep whenever possible, and emptied into a bucket containing seawater. Usually large seagrass clumps encountered were collected randomly, and small fragments were ignored. In the laboratory, the samples were thoroughly rinsed in water and species were identified and counted. *Z. marina* frond lengths were measured from the bottom of the samples to the top of the longest branches (± 1 cm) using a 30 cm ruler.

## Results and discussion

A large area of floating *Z. marina* clumps was observed in our survey (marked in red, Fig. 1.). The area between N 38°58.9358’ E 118°50.1150’ and ‘Point 14’ had frequent clumps, and the area between ‘Point 14’ and ‘Point 15’ had large amounts of floating clumps. In the latter transect line, we estimated the clumps to make up an area of approximately 80–100 m. The Caofeidian seagrass meadows are located approximately 6,400 m away from Point 14, 10,000 m away from Point 15 and 13,500 m away from N 38°58.9358’ E 118°50.1150’. We observed ~10 km^2^ seagrass meadows and large scale areas of floating *Z. marina* clumps around an artificial oil platform in Caofeidian sea areas, near Long island (Fig. 3) in June 2018. Thus, we suggested that these floating seagrass clumps originated from the Caofeidian seagrass meadows. Natural disturbances such as weather, tides and the degree of bed exposure, as well as anthropogenic impacts such as dredging, fishing and anchoring can destroy seagrass meadows and displace plants from the beds to distant locations offshore [8-9, 16-19]. Dierssen et al. [11] found that strong southerly winter winds in Greater Florida Bay advected considerable amounts of seagrass wracks comprised predominantly of *S. filiforme* from the dense meadows in the area to oligotrophic Atlantic Ocean Waters. In our study, isolated *Z. marina* leaves became more aggregated into patches between ‘Point 14’ and ‘Point 15’ and could be found in long windrows produced by downwelling lobes of Langmuir circulation. Aggregations of seagrass vegetation are common in shallow waters due to slow counter-rotating vortices at the ocean’s surface known as Langmuir circulation [20].

**Fig. 3.**
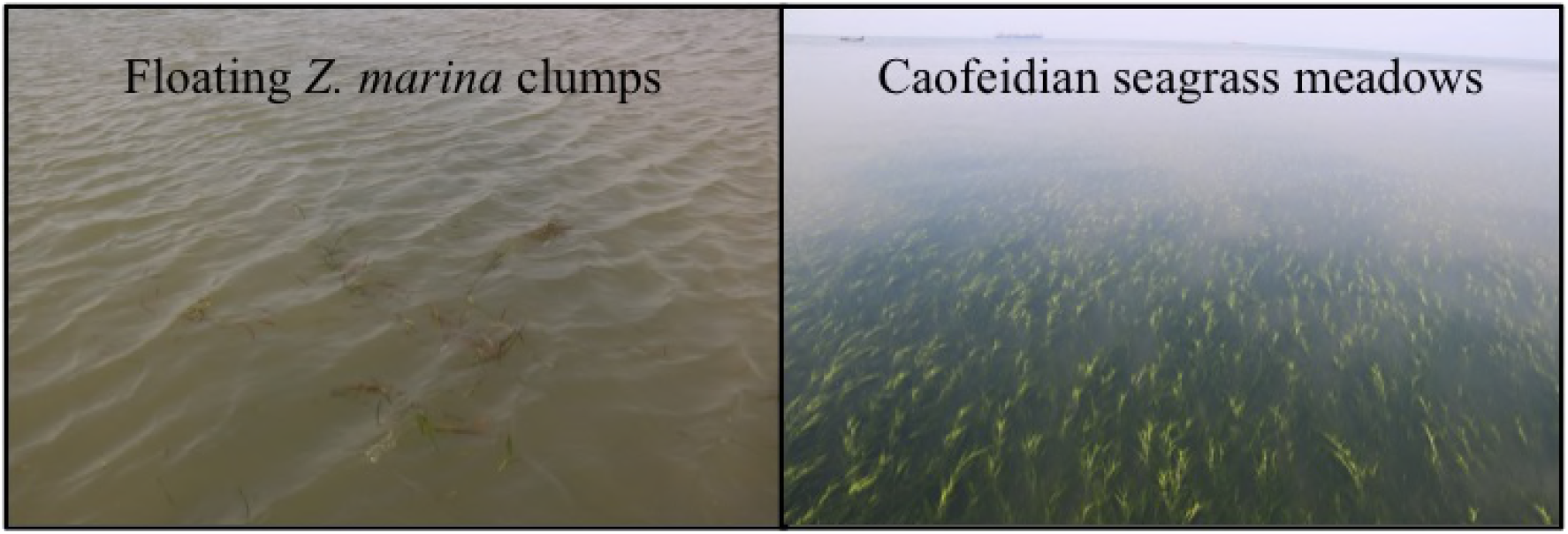
Left: Floating *Zostera marina* clumps found in Caofeidian seagrass meadows; Right: ~10 km^2^ Caofeidian seagrass meadows observed on June 1st 2018.

Generally floating seagrass wracks can be considered a relatively long-lived ecosystem “hot spot”, providing habitats for a wide variety of marine organisms [21]. As floating wracks travel, they release dissolved organic carbon and colored dissolved organic material into surface waters which results in enhanced bacterioplankton in the surrounding seawater. The diverse community supported by floating wracks also serves as a resource for larger pelagic fish, which feed on the smaller organisms living on and near the wracks. However, during the surveys in the present study, marine organisms were not found accompanying the floating seagrass clumps. In addition, through trawling surveys conducted in the sea areas surrounding ‘Point 5’, ‘Point 6’, ‘Point 7’ and ‘Point 8’, we estimated 0.656–1.694 kg/m^2^ (unpublished data) *Z. marina* debris to be laid down in the seabed. Floating seagrass debris provides plentiful nutrition and is an important natural food source for an economically important local aquaculture species, *Apostichopus japonicas*. Song et al. [22] found that the specific growth rates, food utilizing efficiencies and energy budgets of *A. japonicas* were strongly influenced by the ratio of *Z. marina* in their diets.

The dominant frond length category of isolated *Z. marina* leaves in the present study was 40–50 cm in both ‘Line A’ and ‘Line B’ (Fig. 4). In ‘Line A’, > 15% of fronds were in each of the 30–40, 40–50 and 50–60 cm categories. Whereas, in ‘Line B’, > 15% of fronds were in each of the 40–50, 50–60 and 60–70 cm categories. Less than 5% of fronds in both Lines were in the largest size categories of 80–90 or 90–100 cm. In the Caofeidian *Z. marina* seagrass meadows, the canopy height was 15.2 ± 5.84 to 62.1 ± 7.34 cm.

**Fig. 4.**
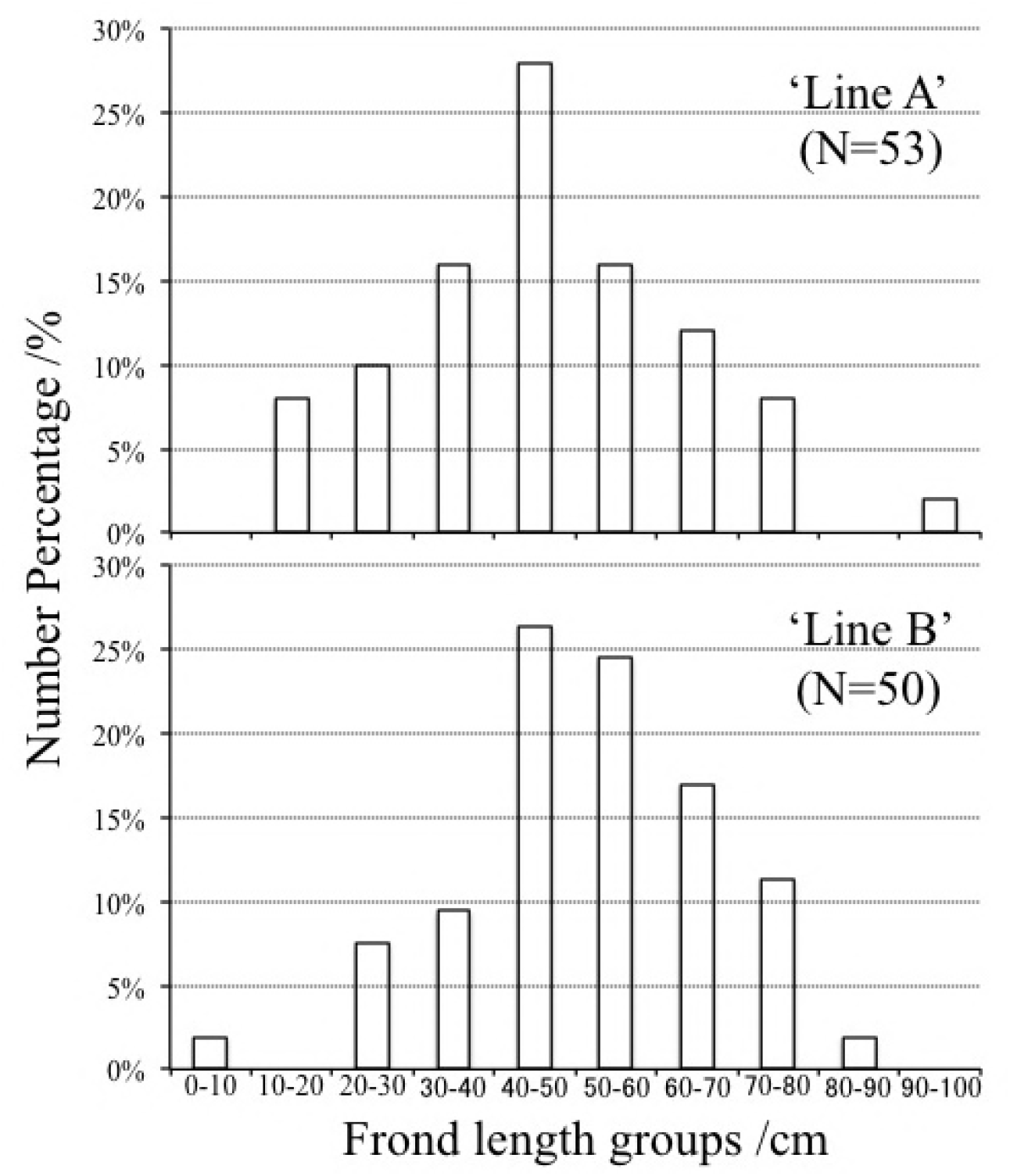
Number percentage (%) of floating *Zostera marina* leaves within different frond length categories (cm) in ‘Line A’ (N = 53) and ‘Line B’ (N = 50).

This study is the first survey to bridge the knowledge gap on floating *Z. marina* clumps in the temperate zones along Chinas coastline. We precluded that the floating seagrass clumps in the northernmost area of Bohai Bay (38°57’1.14”−39° 0’41.28” N, 118°45’23.22”−118°47’6.96”E) originated from the Caofeidian seagrass meadows. Our findings provide a greater understanding about the life cycle of *Z. marina* including how it is transported to the seafloor and becomes organic debris for benthic communities. The dislodgement and break-up characteristics of floating *Z. marina* clumps and the transportation of their organic matter to the seafloor will be the subject of future studies.

## Acknowledgements

The authors thank to Mr Li Chun and the members of Tangshan marine ranching L.t.d. for their helps in field samplings and to members of CAS Key Laboratory of Marine Ecology and Environmental Sciences, Institute of Oceanology, Chinese Academy of Sciences for their constructive discussions and encouragements. The research was supported by Taishan industry leader talent project, the Strategic Priority Research Program of the Chinese Academy of Sciences (Grant No. XDA11020703), Qingdao Postdoctoral Applied Basic research (Grant No. Y7KY02106N), NSFC-Shandong Joint Fund for Marine Science Research Centers (Grant No.U1406403), Key research and development plan (major key technology) project in Shandong (Grant No. 2016ZDJS06A02), National Marine Public Welfare Research Project (Grant No. 201305043), Postdoctoral International Exchange Program Introduction Project (Grant No. Y8KY02102L), Laboratory for Marine Fisheries Science and Food Production Processes and Qingdao National Laboratory for marine science and technology (2016LMFS-B08). This study was financially supported by Creative Team Project of the Laboratory for Marine Ecology and Environmental Science, Qingdao National Laboratory for Marine Science and Technology (NO. LMEES-CTSP-2018-1)

